# Effects of Unexpected Underfoot Perturbations During Turning on Measures of Mediolateral Stability and Corresponding Recovery Strategies

**DOI:** 10.1101/2025.01.30.635610

**Authors:** Tyler K. Ho, Nicholas Kreter, Cameron Jensen, JunSeop Son, Paula Kramer, Peter C. Fino

**Affiliations:** University of Utah, Department of Health & Kinesiology, Salt Lake City, UT; Center for Limb Loss and MoBility, VA Puget Sound Health Care System, Seattle, WA; University of Oregon, Department of Human Physiology, Eugene, OR; University of Nebraska Omaha, Department of Biomechanics, Omaha, NE; University of Utah, Department of Mechanical Engineering, Salt Lake City, UT; University of Utah, Department of Biomedical Engineering, Salt Lake City, UT; University of Utah, Department of Physical Therapy & Athletic Training, Salt Lake City, UT

**Author notes:** Corresponding Author: Peter C. Fino, PhD, 250 S 1850 E, Rm 257, Salt Lake City, UT, 84112, USA.

**Keywords:** Reactive balance, locomotion, margin of stability

## Abstract

Humans regularly walk across uneven terrain, which demands the use of reactive control strategies to maintain forward progress and stability. While reactive control during walking has been well described during straight gait, it is unclear how reactive control differs during turning gait. Because turning is asymmetrical, perturbations to the inside and outside limbs may elicit different reactive adjustments. This study investigates how unexpected underfoot perturbations alter stability measures during turning and how individuals alter their foot placement to maintain stability after such perturbations. Seven healthy adults completed walking trials around a circular track while wearing mechanized shoes that pseudo-randomly delivered underfoot perturbations to either the inside or outside limb. We calculated mediolateral margin of stability corrected for centripetal acceleration (ML MoS*_C_*), step width, and step length from kinematic data. Linear mixed effects models compared the effects of perturbation type (inversion vs eversion), perturbation limb (inside vs outside), and their interaction for each outcome measure. ML MoS*_C_* was affected by both perturbation type and perturbation limb, with larger changes observed during eversion, and outside, perturbations. Changes to step width and step time during two recovery steps after each perturbation were primarily influenced by the perturbation limb – outside perturbations elicited consistent changes during two recovery steps compared to one altered step after inside perturbations. Perturbations to the outside limb during turning disrupt gait longer than perturbations to the inside limb. This difference across perturbation limb may indicate that outside steps are more important to maintaining and recovering stability than inside ones during turning.

## 1. INTRODUCTION

Complex walking environments challenge human stability – the ability to avoid falling – on a daily basis. Walking in such environments demands both anticipatory and reactive control strategies (Hof and Duysens, 2013; Zheng et al., 2011; Bruijn and van Dieën, 2018), with unanticipated balance perturbations necessitating reactive control strategies. These control strategies commonly include changes to foot placement (Bruijn and van Dieën, 2018; Fettrow et al., 2019; Kreter et al., 2022; Reimann et al., 2017), ankle or hip joint torque (Hof et al., 2010), push-off force in the trailing limb (Jin et al., 2023), or step time (Roeles et al., 2018). Such studies examining reactive balance control during straight-line gait have provided insight on effective locomotor control during walking, but fewer studies have examined reactive balance control during non-straight walking, including curved walking and turning.

As many as half of all steps taken throughout a day are non-straight turning steps (Glaister et al., 2007), and turning differs from straight-line gait in several important ways. During turning, the body center of mass (CoM) is biased toward the medial edge of the base of support (Taylor et al., 2005; Xu et al., 2004). This asymmetry is a hallmark of turning (Orendurff et al., 2006), where steps with the outside limb tend to be longer and narrower compared to those with the inside limb. Because turning gait is asymmetrical, perturbations to the inside and outside limbs may affect the CoM trajectory in different ways (Kreter and Fino, 2024). For example, walkers may be more likely to experience a slip on the outside stepping limb compared to on the inside stepping limb (Rasmussen et al., 2022). Asymmetrical ground reaction forces and body inclination angles further suggest that the outside limb is more sensitive to irregular walking surfaces than the inside limb during turning (Yamaguchi et al., 2018, 2012). However, the effects of underfoot perturbations (i.e, uneven walkways or irregular terrain) on turning remain unclear; the majority of uneven walkway tests involve straight walking.

The purpose of this study was to investigate how unexpected underfoot perturbations affect mediolateral (ML) stability during turning gait. Specifically, we aimed to answer three questions: 1) How do unexpected underfoot perturbations affect metrics of ML gait stability during turning? 2) Does the direction of the perturbation (e.g., inversion vs. eversion ankle perturbations) affect the recovery response? 3) Do perturbations to the inside foot elicit different recovery responses than perturbations to the outside foot? To answer these questions, we measured ML margin of stability (MoS) accounting for centripetal acceleration (ML MoS*_C_*), step width, and step time as healthy young adults walked around a circular platform with unexpected, discrete underfoot perturbations. We hypothesized that unexpected underfoot perturbations would decrease ML MoS*_C_* during turning for up to two steps after each perturbation (H1), similar to results seen in straight-line gait (Kreter et al., 2022). We hypothesized that inversion and eversion perturbations would elicit opposite changes in step width and step time on recovery steps following the perturbations (H2). We also hypothesized that perturbations to the outside limb would elicit greater changes to step width and step time than perturbations to the inside limb (H3).

## 2. MATERIALS and METHODS

### 2.1 Participants

Seven healthy young adults [3 female / 4 male, mean (SD) age=23.3 (3.1) years, mean (SD) height=172.9 (8.3) cm, mean (SD) mass=64.74 (9.81) kg] were recruited from the local community for participation in this IRB-approved study. Each participant provided informed written consent prior to participation. Exclusion criteria were: 1) self-reported problems with stability or balance, 2) a history of neuromotor pathology that could impair balance or locomotion, 3) any prior reconstructive surgery to the lower limbs, 4) prescribed or recreational use of psychoactive drugs for 24 hours surrounding data collection, 5) mass >136kg (weight limit of shoe devices used for the perturbations).

### 2.2 Procedures

Participants were fitted with a pair of custom mechanized shoes with two plastic blocks built into the sole of the left shoe such that they would not interfere with normal gait (Figure 1). The plastic blocks were connected to a servo motor, which would rotate them outwards and elicit underfoot perturbations that either inverted or everted the foot (similar to stepping on a small rock). The servo motors were controlled via an Arduino microprocessor and force sensitive resistor that tracked steps. When walking, the Arduino microprocessor would activate the motors randomly every 5-9 strides to deliver the underfoot perturbations. Similar designs have been used in several previous studies that have shown that these small underfoot perturbations are enough to elicit measurable changes in gait (Kim et al., 2013; Kim and Ashton-Miller, 2012; Kreter et al., 2022, 2021).

**Figure 1:**
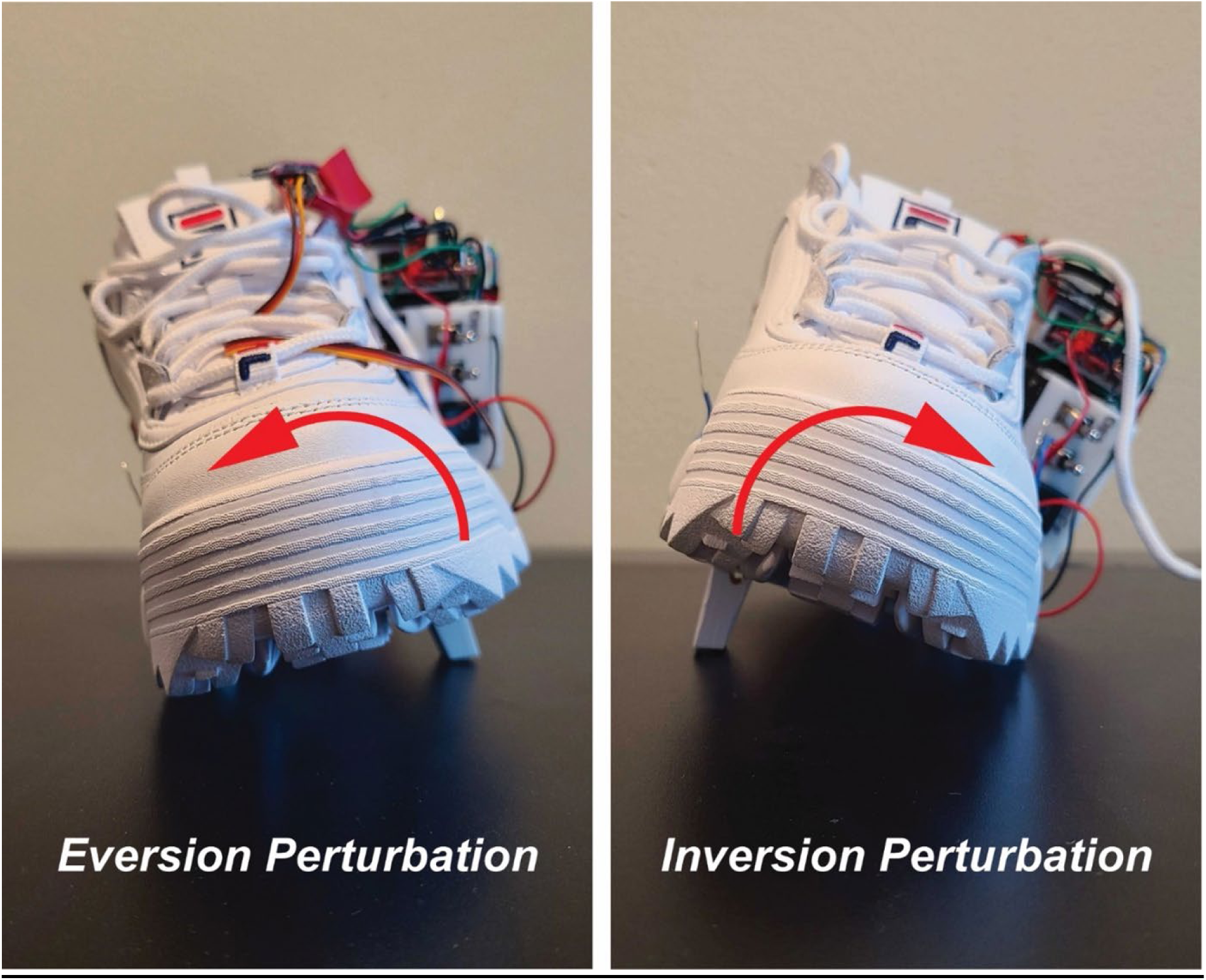
A custom mechanized shoe with two plastic blocks recessed in the sole. Micro-servo motors rotated these blocks out to produce either an eversion (left image) or inversion (right image) perturbation.

All walking trials were completed around a circular track, which had an inner radius of 1.2m, an outer radius of 1.6m, and a thickness of 2.5cm. To mark the progress around the track, the circle was divided into quarter-rings using colored tape. At the start of the session, participants completed a calibration trial to determine their comfortable walking speed around the track. To do so, participants were instructed to walk four laps around the track at their self-selected comfortable walking speed. The average lap time of the middle two laps determined the self-selected walking speed. The first and fourth laps were excluded to prevent influences from gait initiation or termination. Each participant was asked to maintain this comfortable pace throughout testing. Similar to prior studies, we set up a metronome to play a beat once for every one-fourth of their established lap time (Brown et al., 2021; Seddighi et al., 2024). Participants were instructed to complete a quarter lap every metronome beat but were not penalized for missing a mark.

Each participant completed a series of four blocks of trials where they walked either clockwise or counterclockwise and received either inversion or eversion perturbations. Each block started with a 1-minute “acclimation” trial with no perturbations, followed by a three-minute perturbation trial, and concluded with a 1-minute “washout” trial with no perturbations (Figure 2). The order of these blocks was randomized for each participant. Because all perturbations were delivered to the left foot, only the inside foot was perturbed during counterclockwise trials and only the outside foot was perturbed during clockwise trials (Figure 3).

**Figure 2:**
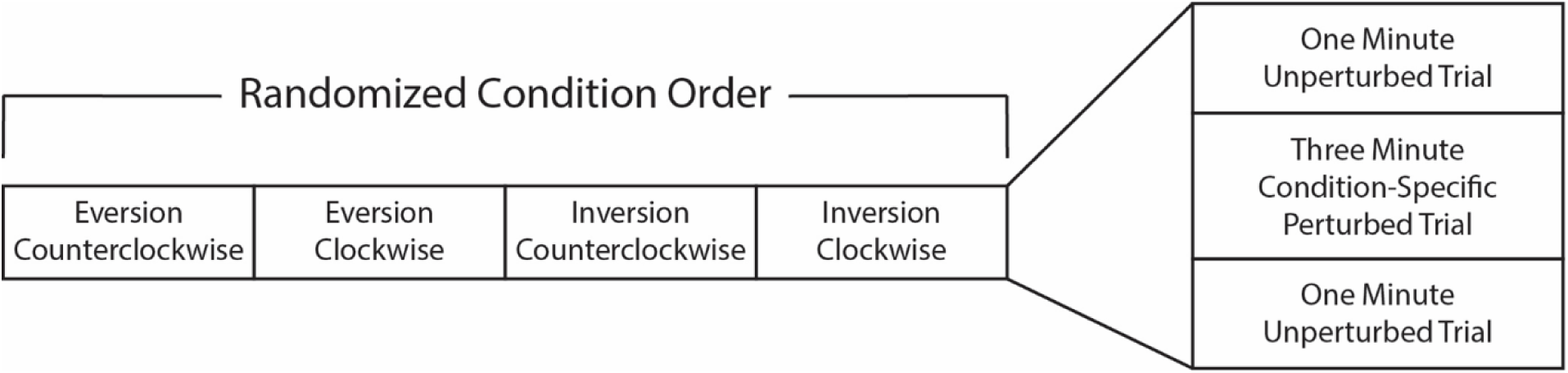
Breakdown of the trial order for this study. During each condition block, participants completed a one minute unperturbed “acclimation” trial, a three minute condition-specific perturbed trial, followed by another one minute “washout” unperturbed trial.

**Figure 3:**
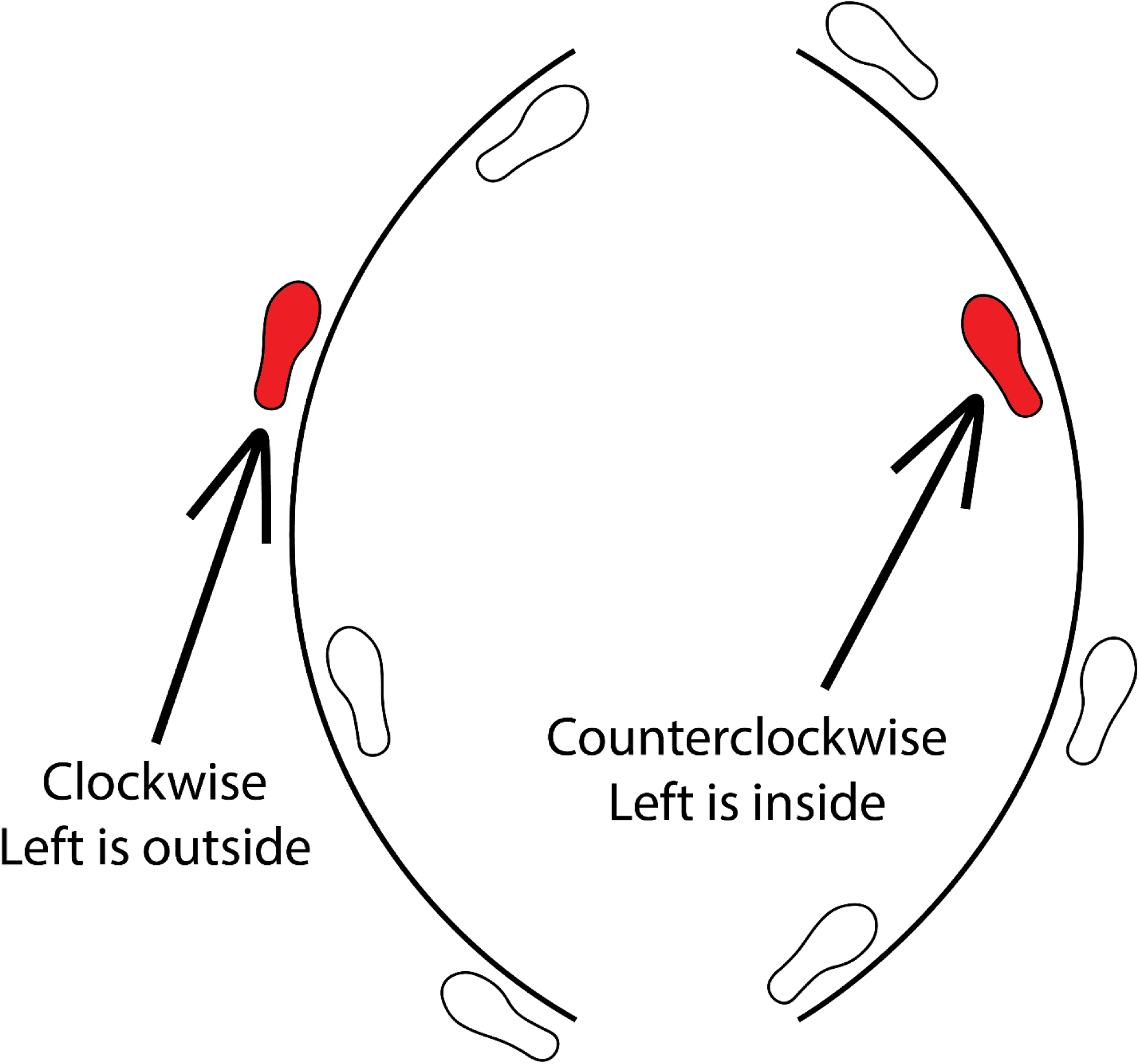
Diagram of left steps as inside vs outside steps during turning. We manipulated the perturbed limb by changing the direction of turning (i.e., clockwise vs counterclockwise).

A 12-camera 3D motion capture system (Nexus 2.12, VICON, Oxford, UK; collected at 200Hz) surrounded the walking environment and captured positional marker data from a custom marker set with 51 retroreflective markers. For our analysis, we used the bilateral retroreflective markers on the pelvis (ASIS and PSIS) and feet (second metatarsophalangeal joint [MTP], fifth MTP and calcaneus) featured in this custom marker set.

### 2.3 Data Analysis

Custom MATLAB codes (ver. R2020b) were used for data processing and data analysis. Positional data were imported and passed through a 4^th^-order phaseless Butterworth filter with a cutoff of 6Hz. The body CoM position was estimated from the four pelvis markers and the positions and orientations of the feet were estimated from the heel, second MTP and fifth MTP markers (Havens et al., 2018). Gait events of heel strike and toe-off were calculated using the positional methods of Ulrich et al. (2019). Steps were separated into inside or outside steps and were categorized as nonperturbed or perturbed steps. The first four steps following a perturbation were categorized as a first, second, third or fourth recovery step (Kreter et al., 2022). A rotating reference frame was aligned with the walking track to define the anteroposterior (AP) and mediolateral (ML) directions (Ho et al., 2023). During each step, the ML direction was determined by the vector from the center of the circular track to the position of the body CoM at contralateral toe-off. The AP aspect was calculated as the cross-product of the ML aspect and a vertical vector, with the positive direction in the direction of travel (Ho et al., 2023).

To quantify the effect of the perturbations on stability, ML margin of stability (MoS) was calculated as the distance between the extrapolated CoM (xCoM) and the lateral edge of the base of support (BoS) (Equation 1) at the moment of contralateral toe-off (Curtze et al., 2024; Hof, 2008; Watson et al., 2021). The typical MoS equation assumes simple inverted pendulum kinematics during gait where the unstable equilibrium point occurs when the body CoM is located directly over the BoS (Hof et al., 2005). However, turning at a constant radius and constant speed shifts this unstable equilibrium point away from vertical to account for centripetal force applied at the foot (Fino et al., 2016). To account for this shift, we applied a correction term based on the velocity of the CoM (*v*), gravity (*g*), the radius of the CoM (*r_CoM_*), and the height of the CoM (*L*) (Equation 2; See Supplement for derivation). Throughout all calculations, a negative MoS or MoS_C_ indicated the participant would fall laterally over the unstable equilibrium point, where a positive MoS_C_ indicated the participant was instantaneously ‘stable’ and would eventually fall back towards the contralateral limb.

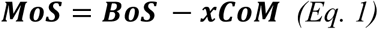

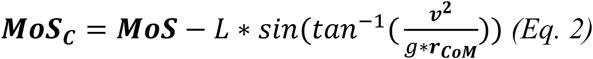

Step width was calculated as the ML aspect of the vector connecting consecutive heel strikes, with the positive direction pointed medially from the initial heel strike in each step cycle (Ho et al., 2023). Step time was calculated as the time between consecutive heel strikes. The means of each measure (ML MoS*_C_*, step width, and step time) for inside and outside steps were calculated across all the normal turning steps (i.e. not perturbed or recovery steps, defined as the perturbation step + 4 recovery steps) within each trial. For each perturbation, we then calculated the change in ML MoS*_C_*, the change in step width, and the change in step time relative to unperturbed steps of the same walking trial.

### 2.4 Statistical Analysis

To test our three hypotheses, we fit linear mixed effects models to our co-primary outcomes: the change in ML MoS*_C_*, the change in step width, and the change in step time relative to unperturbed steps. Linear mixed effects models were fit separately for the perturbed step, first recovery step, and second recovery step. Each model included fixed effects of perturbation type (inversion vs. eversion) and perturbation limb (inside vs. outside), as well as the type*limb interaction. Random intercepts for each participant accounted for repeated measures within subjects. We used pairwise contrasts to test for differences from zero within each condition (i.e., was the perturbed / recovery step different from normal walking; H1), as well as differences across perturbation type (H2) and perturbation limb (H3). All analyses were completed in the Statistics and Machine Learning Toolbox in MATLAB R2020b. A significance level of 0.05 was used throughout.

## 3. RESULTS

### 3.1 Perturbation Step

Main effects of the intercept, perturbation limb, and perturbation type indicated that unexpected underfoot perturbations affected the ML MoS*_C_* during the perturbed step, but the effect varied depending on the type of perturbation and stance limb (Table 1). The perturbation did not affect the step width or step time of the perturbed step.

**Table 1:**
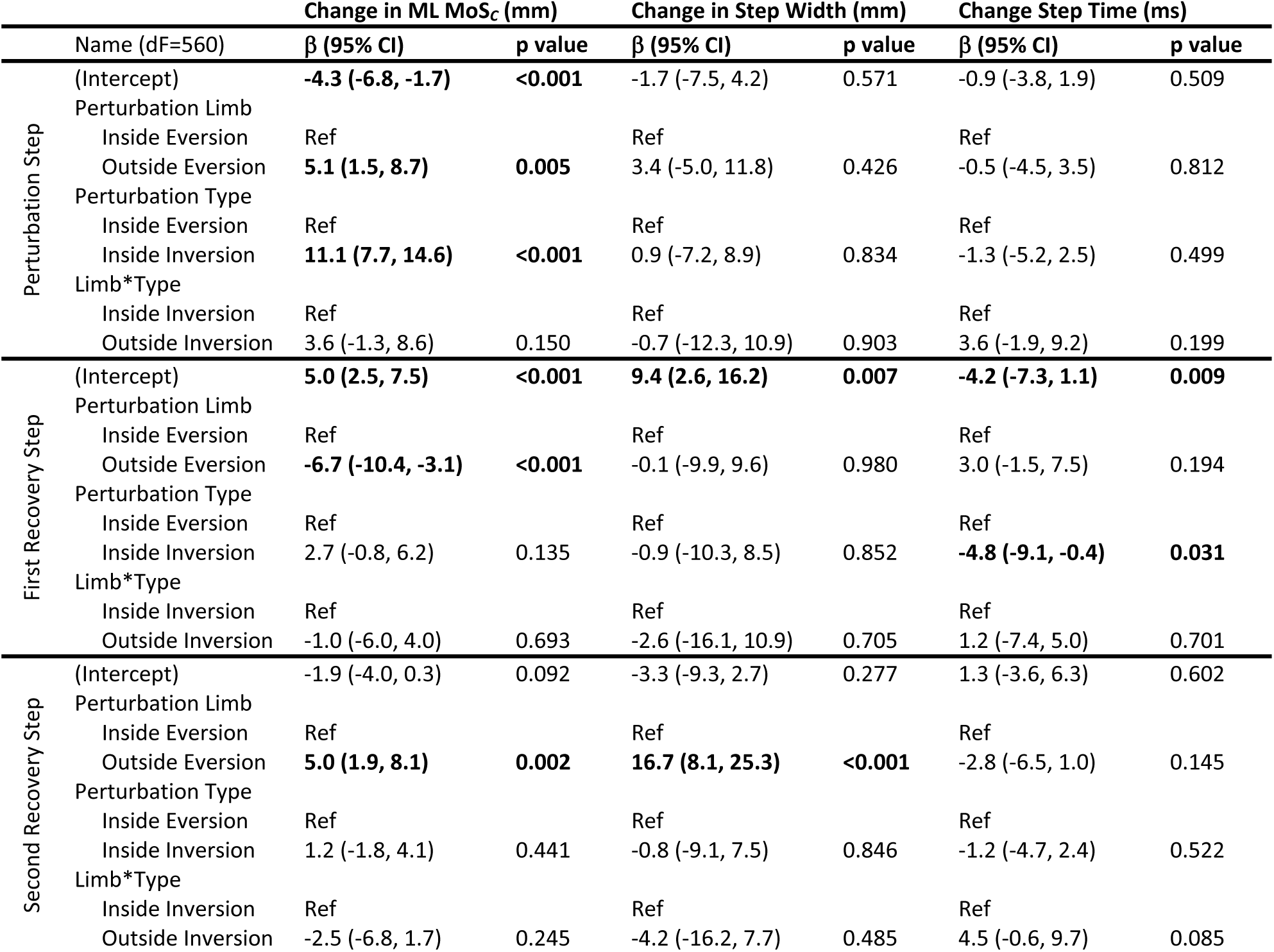
Coefficients and results from the linear mixed models for the change from the mean of ML MoS*_C_*, step width, and step time during each perturbation step and the two subsequent recovery steps. Fixed effects included perturbation limb (inside vs outside), perturbation type (eversion vs inversion) and the interaction of limb*type. The reference condition was the inside eversion perturbation condition. Bolded values indicate statistical significance (α=0.05)

#### 3.1.1 Effect of Perturbation Compared to Normal Gait

Decomposing the linear mixed effects model with contrasts, we observed that ML MoS*_C_* during the perturbed step was significantly smaller than the mean ML MoS*_C_* of unperturbed steps during inside eversion perturbations (Beta = −4.3mm, *p*=0.001), but greater than normal walking during inside inversion perturbations (Beta = 6.8mm, *p*<0.001). Though ML MoS*_C_* did not differ from unperturbed gait during outside eversion perturbations (Beta = 0.8mm, *p*=0.514), ML MoS*_C_* was greater during outside inversion perturbations (Beta = 15.5mm, *p*<0.001).

#### 3.1.2 Effect of Perturbation Type and Limb

There was a significant difference in ML MoS*_C_* between perturbation types across both inside (*p*<0.001) and outside limb perturbations (*p*<0.001). There was also a significant difference in ML MoS*_C_* between perturbation limbs across both eversion (*p*=0.005) and inversion perturbations (*p*<0.001).

### 3.2 First Recovery Step

Step width, step time, and ML MoS*_C_* of the first recovery step all differed from unperturbed walking for at least one condition (Table 1).

#### 3.2.1 Effect of Perturbation Compared to Normal Gait

Decomposing the linear mixed effects model using contrasts, we observed that participants took a wider first recovery step after inside eversion (Beta = 9.4mm, *p*=0.007), inside inversion (Beta = 8.5mm, *p*=0.010), and outside eversion perturbations (Beta = 9.3mm, *p*=0.009), but not after outside inversion perturbations (Beta = 5.8mm, *p*=0.088). Participants also took a faster first recovery step after inside eversion (Beta = −4.2ms, *p*=0.009), inside inversion (Beta = −9.0ms, *p*<0.001), and outside inversion perturbations (Beta = −4.8ms, *p*<0.001), but not after outside eversion perturbations (Beta = −1.2ms, *p* = 0.453). ML MoS*_C_* also remained elevated compared to normal walking after inside limb perturbations (inside eversion Beta = 5.0mm, *p* < 0.001; inside inversion Beta = 7.7mm, *p*<0.001), but not after outside limb perturbations (outside eversion Beta = −1.7mm, *p*=0.192; outside inversion Beta = 0.0mm, *p*=0.952).

#### 3.2.2 Effect of Perturbation Type and Limb

We observed a significant difference in the first recovery step time according to perturbation type across both inside (*p*=0.031) and outside limb perturbations (*p*=0.009). ML MoS*_C_* during the first recovery step differed according to perturbation limb across both eversion (*p*<0.001) and inversion perturbations (*p*<0.001).

### 3.3 Second Recovery Step

Step width and ML MoS*_C_* of the second recovery step both differed from unperturbed walking for at least one outside limb perturbation condition (Table 1).

#### 3.3.1 Effect of Perturbation Compared to Normal Gait

Decomposing the linear mixed effects model using contrasts, we observed that participants took a wider second recovery step after outside limb perturbations, regardless of perturbation type (outside eversion Beta =13.4mm, *p*<0.001; outside inversion Beta = 8.4mm, *p*=0.006), but exhibited no differences in step width two steps after inside limb perturbations. ML MoS*_C_* was also elevated during the second recovery step after outside eversion perturbations (Beta = 3.1 mm, *p*=0.006), but returned to normal across all other conditions.

#### 3.3.2 Effect of Perturbation Type and Limb

We observed a difference in second recovery step width according to perturbation limb across eversion (*p*<0.001) and inversion perturbations (*p*=0.003). There was also a difference in ML MoS*_C_* during the second recovery step according to perturbation limb across eversion perturbations (*p*=0.002).

## 4. DISCUSSION

This study aimed to investigate the effect of unexpected underfoot perturbations on stability during turning. Our three hypotheses were that: 1) participants’ ML MoS*_C_* would deviate from normal walking during perturbed steps and up to two recovery steps, 2) participants would make opposite changes in their step width and step time in response to inversion and eversion perturbations, and 3) participants would make larger changes in their step width and step time in response to outside limb perturbations compared to inside limb perturbations. We found that ML MoS*_C_* significantly changed during all perturbed steps except eversion perturbations to the outside limb. ML MoS*_C_* was greater, compared to unperturbed turning gait, during the first recovery step after inside limb perturbations, and during the second recovery step after an outside eversion perturbation. We also found that outside limb perturbations elicited more changes in step width and step time than inside limb perturbations. The only consistent difference between perturbation types (i.e., inversion vs eversion) across limbs was observed in the step time of the first recovery step; participants took a faster step after an inversion perturbation than after an eversion perturbation, regardless of the perturbed limb. These results suggest that perturbations during turning create different effects based on the type of perturbation and the stance limb that is perturbed, and that contributions to balance recovery during turning may be asymmetrical.

Changes in ML MoS*_C_* suggest perturbations during turning can differentially affect balance control and forward progression along a desired path. ML MoS*_C_* decreased during inside eversion perturbations, but increased during inversion perturbations. However, such increases do not necessarily mean participants were more stable during an inversion underfoot perturbation. A more positive ML MoS*_C_* here indicates medial instability (Curtze et al., 2024), which also necessitates recovery strategies to re-establish normal walking. Importantly, the effect of medial instability during turning differs by inside and outside limb (see Figures 4-6); medial instability for the outside limb indicates instability *towards* the center of the turn, while medial instability for the inside limb indicates instability *away from* the center of the turn. With this perspective, the change in ML MoS*_C_* during inside eversion perturbations and during outside inversion perturbations both indicate instability towards the center of the circle. In contrast, the change in ML MoS*_C_* during inside inversion perturbations indicates instability away from the center of the circle. Therefore, inversion perturbations consistently elicit changes in ML MoS*_C_* towards the contralateral limb, but these changes have different effects on CoM progression around a curve based on whether the inside or outside limb is perturbed.

**Figure 4:**
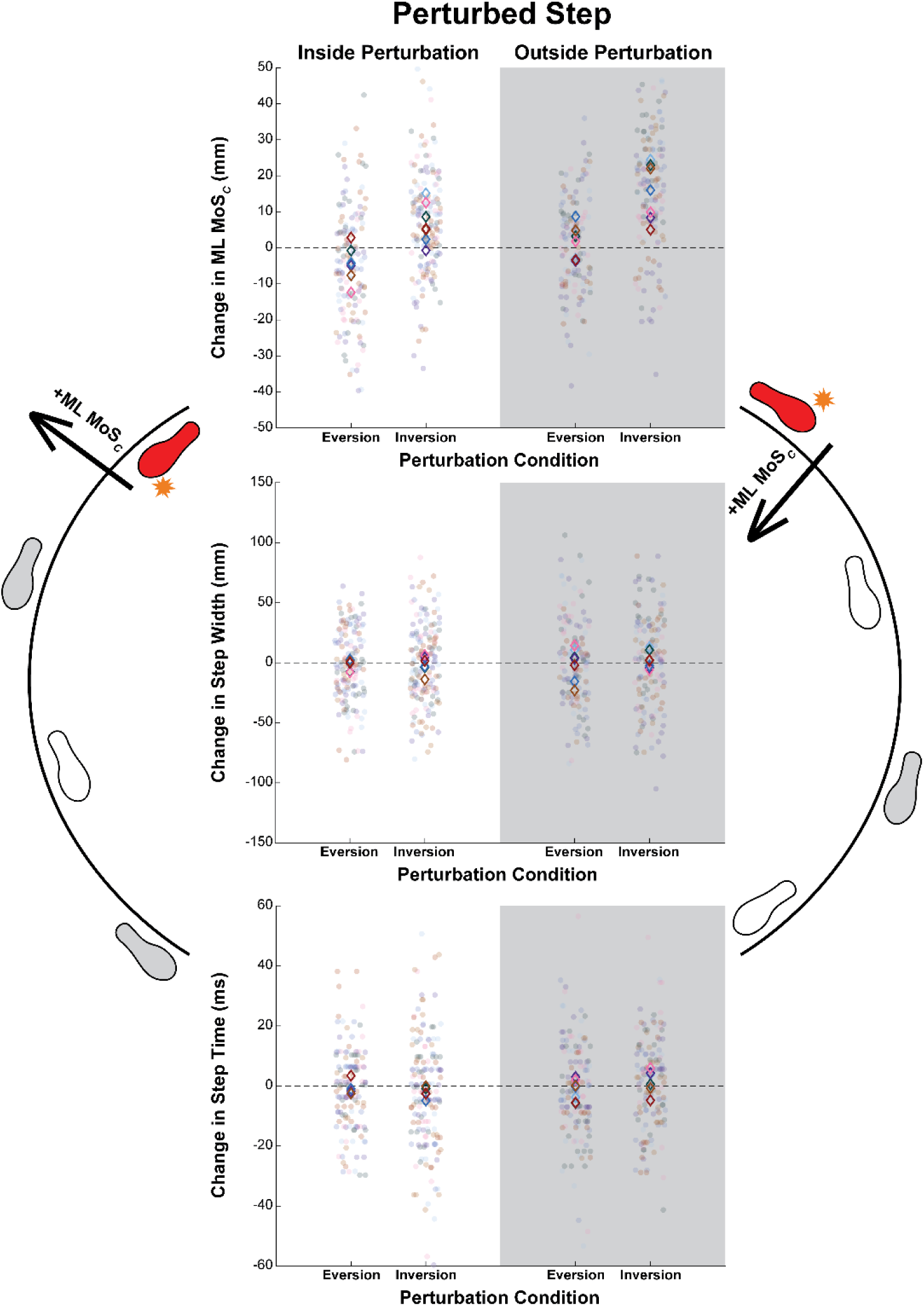
Results of each outcome (the change in ML MoS*_C_*, step width and step time) during each perturbed step. These results were separated by eversion vs inversion perturbations, and by inside limb (white) vs outside (grey) limb perturbations. Each dot represents a single step and the open diamonds represent the mean for each participant. The diamonds and dots are color coded per participant.

**Figure 5:**
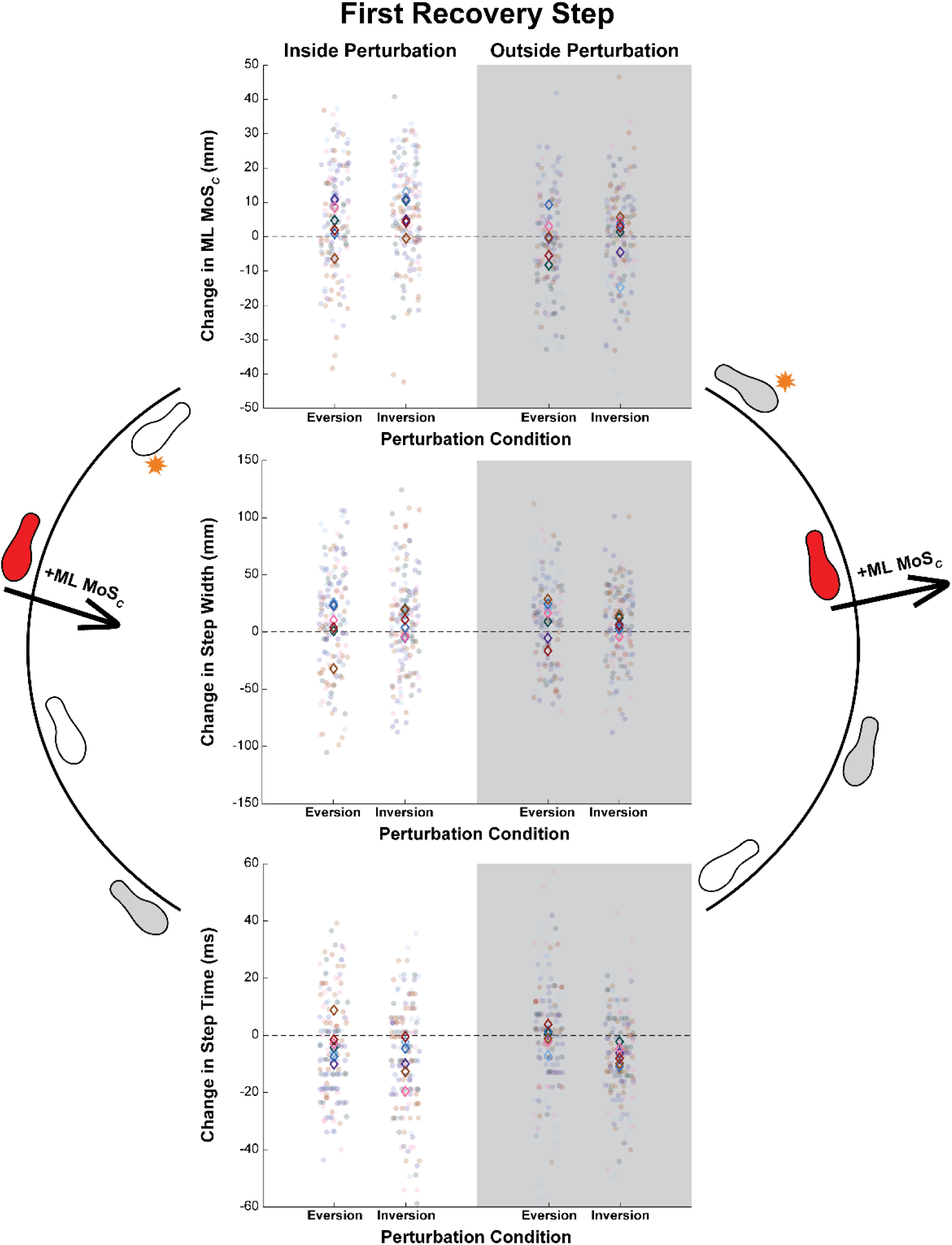
Results of each outcome (the change in ML MoS*_C_*, step width and step time) during each first recovery step. These results were separated by eversion vs inversion perturbations, and by inside limb (white) vs outside (grey) limb perturbations. Each dot represents a single step and the open diamonds represent the mean for each participant. The diamonds and dots are color

**Figure 6:**
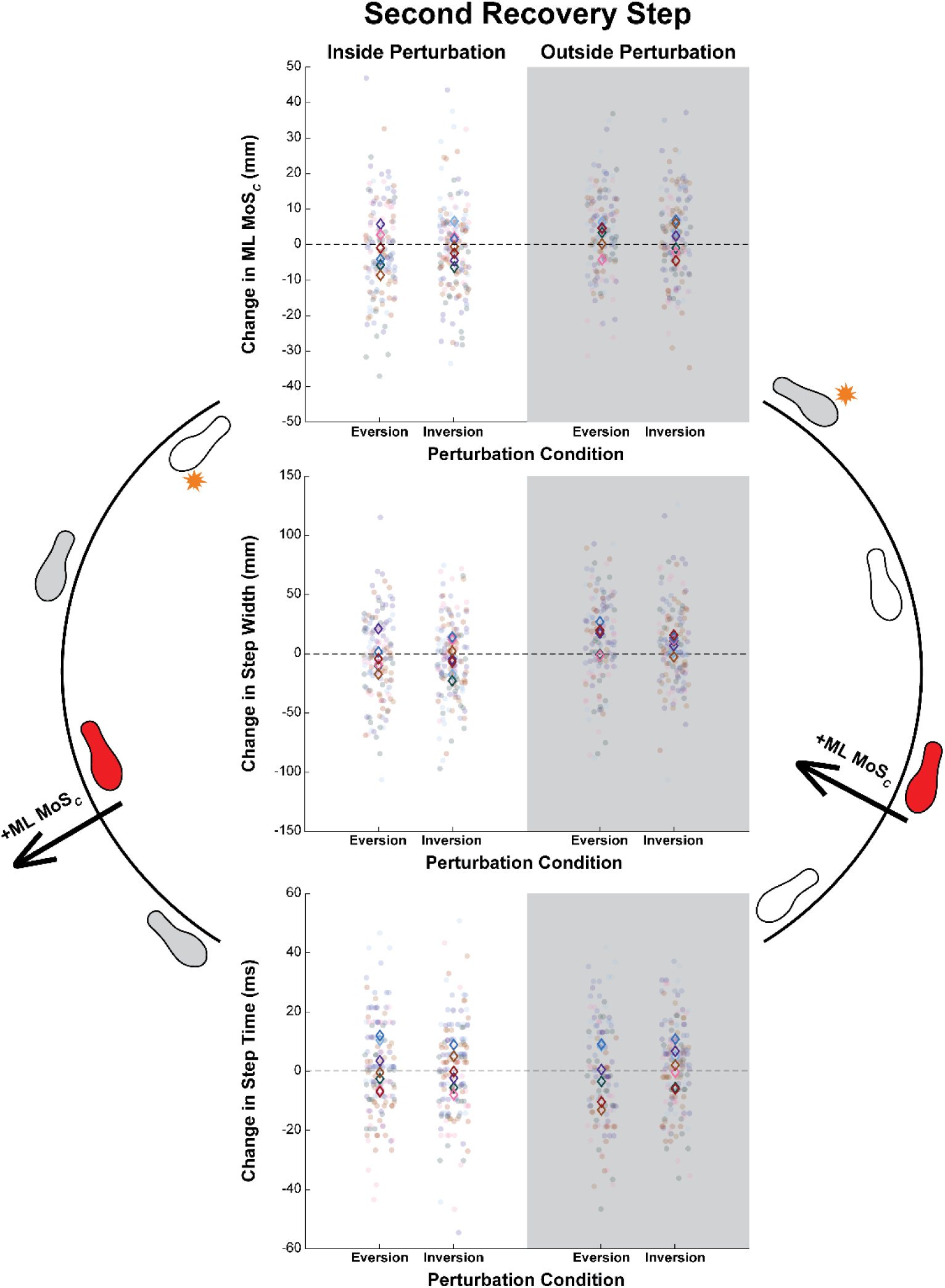
Results of each outcome (the change in ML MoS*_C_*, step width and step time) during each second recovery step. These results were separated by eversion vs inversion perturbations, and by inside limb (white) vs outside (grey) limb perturbations. Each dot represents a single step and the open diamonds represent the mean for each participant. The diamonds and dots are color

Regardless of the effect limb laterality or perturbation type, participants did not return to normal gait until they could make a corrective step with their outside limb. Participants took wider and faster first recovery steps after perturbations to the inside limb (i.e., where the first recovery step was with the outside limb). These corrections were sufficient to return to normal gait at the second recovery step. In contrast, participants took two steps to recover from perturbations to the outside limb, but the specific recovery strategy depended on perturbation type. That is, during the first recovery step following a perturbation to the outside limb (i.e., where the first recovery step was with the inside limb), participants exhibited wider recovery steps for outside eversion perturbations and faster recovery steps for outside inversion perturbations. Further, participants continued to exhibit a wider second step after perturbations to the outside limb (i.e., where the second recovery step was with the outside limb).

The different recovery strategies between perturbations to inside and outside limbs may also be explained as an attempt to limit the curvature of the CoM trajectory. Assuming that the optimal instantaneous CoM trajectory is tangent to the curvature of the path, a perturbation that produces instability away from the center of the circle creates a new CoM trajectory that will never intercept the circle unless corrected (Figure 8). Immediately correcting this perturbation may be most effective to limit the CoM’s divergence away from the circular path. Alternatively, perturbations that produce instability toward the center of the circle create a new CoM trajectory that will eventually re-intercept the circle (Figure 8). So long as the perturbation does not pose an immediate threat of falling, someone may choose to correct the CoM trajectory on either the first or second recovery step. Here, participants made adjustments to step width throughout the first outside recovery step (i.e., one step after perturbations to the inside limb, or two recovery steps after perturbations to the outside limb). Overall, these results suggests that adjustments to outside steps may be preferred over adjustments to inside steps when recovering stability during turning (Rasmussen et al., 2022).

**Figure 7:**
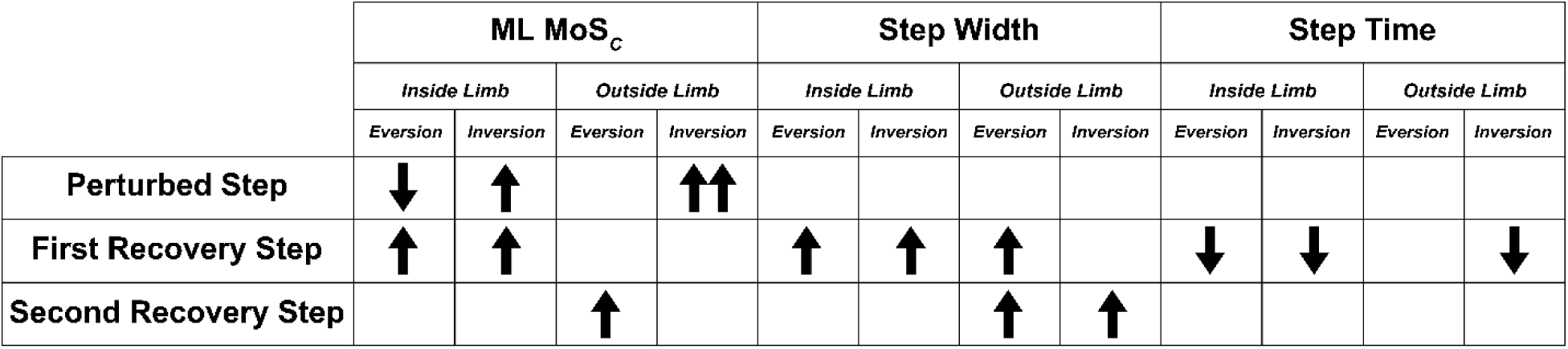
Descriptive table of the direction of change (i.e., increase or decrease) of each outcome (ML MoS*_C_*, step width and step time) in response to each perturbation condition. Blank spaces indicate no significant change from the mean, while an arrow pointing up or down indicates an increase or a decrease respectively. A double arrow indicates a change in one condition that is significantly greater than changes in other conditions.

**Figure 8:**
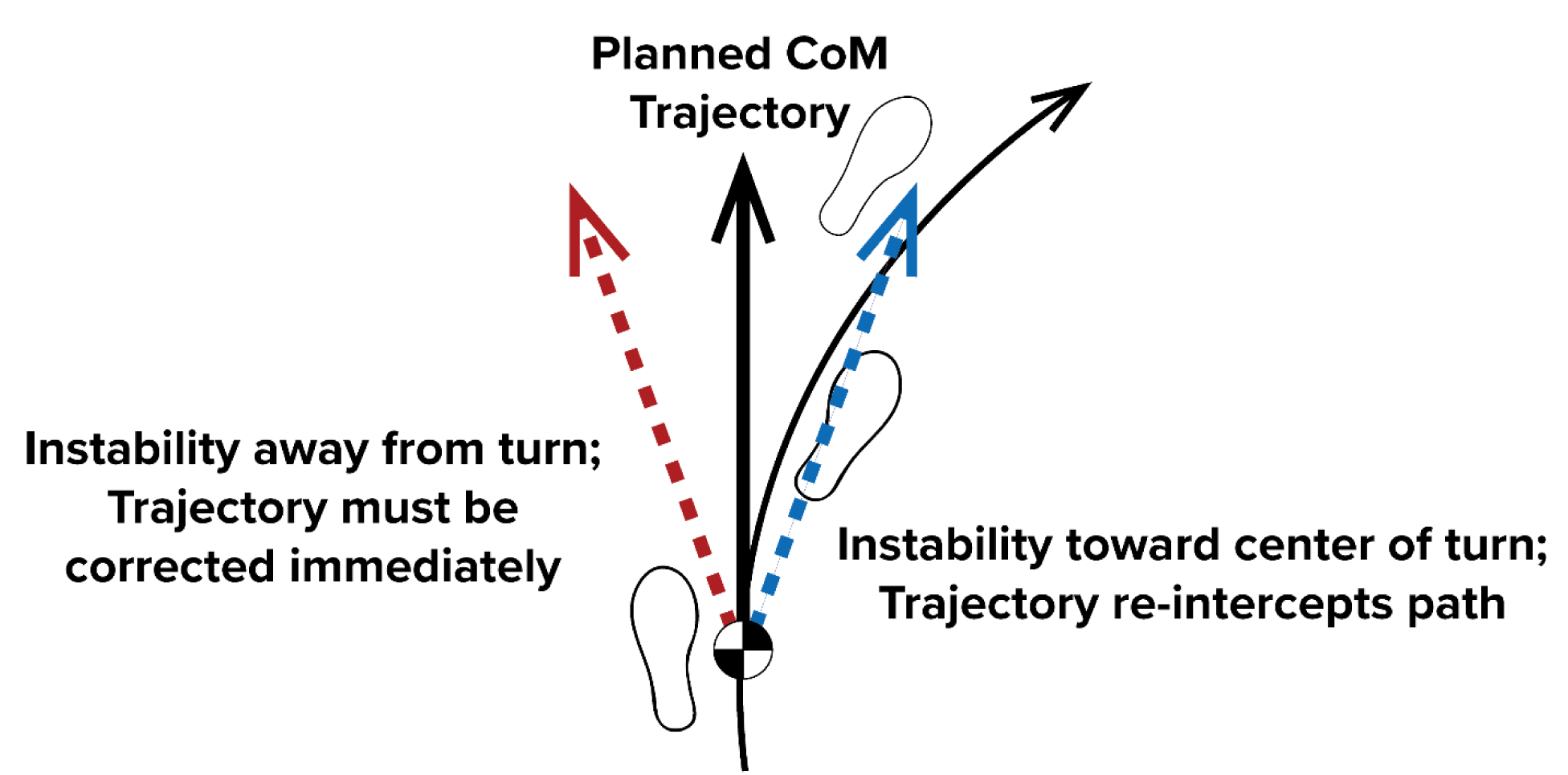
Illustration describing the change in CoM trajectory from instability away from the direction of the turn (red) and instability toward the center of the turn (blue). The altered trajectory away from the turn (red) will continue to diverge from the planned turning path unless corrected, while the altered trajectory toward the center of the turn can be corrected anytime across the first two recovery steps. In this case, walkers will likely opt for the recovery strategy that minimizes the energy required.

Alternatively, limitations of the circular walkway may explain this delay in recovery after an outside limb perturbation. We found that participants tended to be biased toward the inside border of the track, and as such, they may not have had enough space to make the step width adjustment that they normally would have in an uncontrolled environment. This interpretation seems unlikely, given that similar results report bias towards recovering on the outside limb during unconstrained turning (Kreter and Fino, 2024; Rasmussen et al., 2022) and that humans tend to be biased against trajectories that force sharper turns based on energetic demands (Brown et al., 2021).

Notably, the changes in ML MoS_C_, step width, and step time elicited by the perturbation are small (i.e., on the scale of millimeters and milliseconds). Despite the small magnitudes, the changes in ML MoS*_C_* and step width are of a comparable magnitude as seen in previous studies that have used similar devices to deliver underfoot perturbations (Kim and Ashton-Miller, 2012; Kreter et al., 2022), which have shown clinical significance in predicting falls (Allet et al., 2014). We suspect such differences would extrapolate to larger magnitude perturbations that require larger changes in gait, but future research should consider different types (e.g., at different locations along the foot, or different mechanisms) and magnitudes of perturbations.

## 5. CONCLUSION

Underfoot perturbations disrupted ML MoS*_C_* differently across perturbation type, and the change in ML MoS*_C_* was dependent on the limb being perturbed. After these underfoot perturbations during turning, participants employed similar recovery strategies as previously seen in linear walking (i.e., increased step width and decreased step time) (Kim et al., 2013; Kreter et al., 2022). However, the magnitude of these changes in foot placement and timing was also largely determined by the perturbed limb, as participants tended to wait until an outside limb recovery step to correct the ML MoS_C_. These results demonstrate that outside steps may be more important for maintaining and recovering stability during turning compared to inside steps, which may be used more for maintenance of the status quo rather than corrective control.

## ACKNOWLEDGEMENTS

Special thanks to Claire Rogers for her work in developing the data analysis for this study, and to Jasmine Arreguin for her assistance with data processing.

