## Supplement for "Effects of Unexpected Underfoot Perturbations During Turning on Measures of Mediolateral Stability and Corresponding Recovery Strategies"

Supplemental Materials - *Derivation of corrected Margin of Stability formula to account for centripetal acceleration during turning*

The mediolateral Margin of Stability (*MoS*) is traditionally calculated as the distance between the vertical projections of the base of support (*BoS*) and the extrapolated center of mass (*xCoM*; Curtze et al., 2024). The following formula is simplified from the model originally proposed by Hof et al. (2005):

$\boldsymbol{MoS}\mathbf{=}\boldsymbol{BoS}\mathbf{-}\boldsymbol{xCoM}$ *(Eq. S1)*

Equation S1 assumes that the stable equilibrium point occurs when the *xCoM* is directly over the top of the BoS (i.e., MoS = 0). However, during a turn, the CoM experiences centripetal acceleration, which effectively shifts the equilibrium point away from vertical. So, we must add a new term (*C*) to Equation A describe this shift from centripetal acceleration, such that the equilibrium point will still be achieved when *MoS* = 0.

$\boldsymbol{MoS}\mathbf{=}\boldsymbol{BoS}\mathbf{-}\boldsymbol{xCoM-C}$ *(Eq. S2)*

Within the mediolateral plane, *C* reflects the displacement of the center of mass necessary to balance the moments in the frontal plane due to the horizontal ground reaction force and the vertical ground reaction force (Fino et al., 2016)

$\boldsymbol{M}_{\boldsymbol{C}}\boldsymbol{=}\boldsymbol{M}_{\boldsymbol{g}}$ *(Eq. S3)*

These moments are produced by the force due to friction (*F_f_*) and the normal force (*F_N_*) acting on the *BoS*. The moment arm between the foot contact and the center of mass can be estimated by the mean height of the center of mass (*L*) and the frontal plane inclination angle *θ*.

$\boldsymbol{F}_{\boldsymbol{f}}\boldsymbol{L*cos(\theta)=}\boldsymbol{F}_{\boldsymbol{N}}\boldsymbol{L*sin(\theta)}$ *(Eq. S4)*

*F_f_* can be described by the coefficient of friction (*μ*), mass (*m*) and gravity (*g*), while *F_N_* is proportional to *m* and *g*.

$\boldsymbol{\mu mg*cos(\theta)=mg*sin(\theta)}$ *(Eq. S5)*

$\boldsymbol{\mu*cos(\theta)=sin(\theta)}$ *(Eq. S6)*

$\boldsymbol{\mu=tan(\theta)}$ *(Eq. S7)*

Here, the *F_f_* is the force that facilitates the centripetal acceleration in the transverse plane. Therefore

$\boldsymbol{F}_{\boldsymbol{f}}\boldsymbol{=m}\boldsymbol{a}_{\boldsymbol{c}}\boldsymbol{=}\frac{\boldsymbol{mv}^{\boldsymbol{2}}}{\boldsymbol{R}}\boldsymbol{=}\boldsymbol{\mu mg}$	*(Eq. S8)*

$\boldsymbol{\mu=}\frac{\boldsymbol{v}^{\boldsymbol{2}}}{\boldsymbol{gR}}$	*(Eq. S9)*

Where *R* is the radius between the center of mass and the center of the turn and *v* is the tangential velocity of the center of mass. Thus, *θ* can be described as

$\boldsymbol{\theta=}\boldsymbol{tan}^{\boldsymbol{-1}}\boldsymbol{(}\frac{\boldsymbol{v}^{\boldsymbol{2}}}{\boldsymbol{gR}}\boldsymbol{)}$	*(Eq. S10)*

Combining Eqs. S4 and S10, the distance between the *BoS* and the position of the CoM yields

$\boldsymbol{C=L*sin(}\boldsymbol{tan}^{\boldsymbol{-1}}\left( \frac{\boldsymbol{v}^{\boldsymbol{2}}}{\boldsymbol{gR}} \right)\boldsymbol{)}$	*(Eq. S11)*

and substituting this into Eq. S2 yields

$\boldsymbol{MoS}_{\boldsymbol{C}}\mathbf{=}\boldsymbol{MoS}\mathbf{-}L\mathbf{*}sin \boldsymbol{(}{tan}^{-1}\boldsymbol{(} \frac{\boldsymbol{v}^{\boldsymbol{2}}}{g\boldsymbol{*}\boldsymbol{r}_{\boldsymbol{CoM}}}\boldsymbol{)}\boldsymbol{)}$ *(Eq. S12)*

Eq. S12 could also be further simplified

$\boldsymbol{MoS}_{\boldsymbol{C}}\mathbf{=}\boldsymbol{MoS}\mathbf{-}L\mathbf{*}\frac{\frac{\boldsymbol{v}^{\boldsymbol{2}}}{g\boldsymbol{*}\boldsymbol{r}_{\boldsymbol{CoM}}}}{\sqrt{\boldsymbol{1+(}{\frac{\boldsymbol{v}^{\boldsymbol{2}}}{g\boldsymbol{*}\boldsymbol{r}_{\boldsymbol{CoM}}}\boldsymbol{)}}^{\boldsymbol{2}}}}$ *(Eq. S13)*
